# Dynamic DNA methylation changes in the *COMT* gene promoter region in response to mental stress and its modulation by transcranial direct current stimulation

**DOI:** 10.1101/2021.10.01.462774

**Authors:** Ariane Wiegand, Arne Blickle, Christof Brückmann, Simone Weller, Vanessa Nieratschker, Christian Plewnia

**Author notes:** **Corresponding author:** Ariane Wiegand, M.Sc.; Department of Psychiatry and Psychotherapy; Molecular Psychiatry; University of Tübingen, Calwerstraße 14, D-72076 Tübingen, Germany. Contributed equally (Vanessa Nieratschker, Christian Plewnia).

## Abstract

Changes in epigenetic modifications present a mechanism how environmental factors like the experience of stress can alter gene regulation. While stress-related disorders have consistently been associated with differential DNA methylation, little is known about the time scale in which these alterations emerge. We investigated dynamic DNA methylation changes in whole blood of 42 healthy male individuals in response to a stressful cognitive task, its association with concentration changes in cortisol and its modulation by transcranial direct current stimulation (tDCS). We observed a continuous increase in *COMT* promotor DNA methylation which correlated with higher saliva cortisol levels and was still detectable one week later. However, this lasting effect was suppressed by concurrent activity-enhancing anodal tDCS to the dorsolateral prefrontal cortex. Our findings support the significance of gene-specific DNA methylation in whole blood as potential biomarkers for stress-related disorders. Moreover, they suggest alternative molecular mechanisms possibly involved in lasting behavioral effects of tDCS.

## Introduction

Epigenetic patterns are known to be dynamic and associated with environmental factors. Without altering the DNA sequence, epigenetic modifications affect chromatin structure and gene expression. Currently, one of the best studied epigenetic modifications is DNA methylation (DNAm), which plays an important role in gene regulation ^1^. One factor which has consistently been associated with differential DNAm, is stress ^2,3^. Many studies link early life stress to long lasting differences in DNAm ^4,5^ or correlate severe psychiatric symptoms caused by traumatic and stressful life events with differential DNAm profiles ^6^. Hence, epigenetic alterations might be an underlying mechanism how exposure to stress increases the risk of developing psychiatric disorders. Nevertheless, only little is known about the short-term dynamics of methylation changes after stress exposure. Immediate changes in DNAm can be induced by chemical stressors like dimethyl sulfoxide (DMSO) ^7-9^ and can already occur within 20 min after T-cell activation ^10^. Furthermore, an experimental psychological stressor, the Trier Social Stress Test, has shown to be associated with dynamic DNAm alterations in a stress-associated gene within 90 min after exposure ^11^. The dynamic malleability of methylation changes in genes involved in cognitive stress is therefore a potentially critical mechanism for the regulation of human behavior. However, a precise characterization of the degree and time course of these changes is required. Moreover, opportunities to influence this process would be useful and may open new perspectives for individualized therapeutic strategies.

To this aim, we use transcranial direct current stimulation (tDCS), a non-invasive brain stimulation technique, which has been shown to modulate neuroplasticity ^12^. Many studies have demonstrated the impact of tDCS on cognitive processes and training ^13,14^. Most importantly it has been discussed as a potential treatment approach for neuropsychiatric disorders which are often associated with aberrant brain activation patterns ^15^. However, so far little is known about the underlying molecular mechanisms of stimulation effects and how they potentially manifest as long-lasting cognitive improvements and amelioration of psychiatric symptoms. As epigenetic modifications present a mechanism how environmental factors can influence physiological reactions and, moreover, seem to be involved in the pathophysiology of psychiatric disorders they might also be important for the manifestation of tDCS effects.

There is accumulating evidence that genetic factors interact with stimulation effects and contribute to inter-individual variability in tDCS responses ^16^. Particularly, the Val108/158Met polymorphism of the catechol-O-methyltransferase (*COMT*) gene that regulates the dopamine metabolism ^17^ is associated with differential tDCS effects on executive functions ^16,18^. COMT is involved in the degradation of dopamine and, therefore, plays a critical role in cognitive processes and executive functioning ^19,20^. A physiological concentration of dopamine in the prefrontal cortex is important for optimal cognitive functioning ^21^ and a dysregulation of the dopaminergic system is associated with the pathophysiology of neurological and psychiatric disorders like schizophrenia and depression ^22^. Furthermore, it has been shown that acute stress leads to an activation of the dopaminergic system ^23,24^ and the *COMT* gene seems to be crucially involved in the stress response ^25^. The *COMT* genotype, which influences the enzyme’s stability and hence dopaminergic activity ^17^, is also associated with an altered cortisol response ^26,27^. Therefore, the promotor region of this gene appears to be a promising candidate to exemplify the epigenetic signatures of mental stress and its malleability by tDCS.

Thus, the present study aims at *i)* determining the effects of tDCS on task performance, negative affect, and the physiological stress response, *ii)* testing the notion that DNAm of the *COMT* gene is subject to immediate modulation by mental stress, and *iii)* providing initial evidence that tDCS can influence this process.

## Methods

### Participants

The study sample was recruited in two cohorts. As a pilot study, 22 healthy participants (mean age: 23.6 years, SD = 3.0; mean years of formal education: 16.9, SD = 3.3) took part in the experiment. The effects of tDCS on task performance and affect were described in Wiegand et al. (2019) in more detail ^28^. See Supplementary Figures S1 and S2 for *COMT* DNAm and cortisol data from this pilot cohort. Since, to our knowledge, this is the first human study investigating dynamic DNAm in the context of cognitive stress and its modulation by tDCS, no previously reported effect sizes for a power and sample size estimation were available. To increase reliability of our findings, we replicated the same experiment with another 20 healthy participants (mean age: 23.3 years, SD = 3.5; mean years of formal education: 14.4, SD = 6.2). See Supplementary Figures S3 and S4 for *COMT* DNAm and cortisol data from this replication cohort. Since there were no prominent differences between the data from the two cohorts, results in the main manuscript are reported for the merged cohort including all 42 participants. The inclusion criteria, experimental procedure, and sample handling and storage were the same for both cohorts and instructions were given by the same instructor using a detailed script. All participants were recruited within two years. The two cohorts showed no significant differences with respect to age (*t*(40) = -0.39, *p* = 0.97) or years of education (*t*(40) = 1.13, *p* = 0.26). To reduce inter-individual DNAm variability, all participants were aged between 18-30 years, male, non-smoking, and of European descent. Furthermore, screening excluded participants with a history of mental or neurological illness, relevant somatic disorders (two participants were suffering from hypothyroidism), dyscalculia, metallic foreign particles around the head, a cardiac pacemaker, and the usage of psychotropic or other medication that may impact DNAm status (two participants took L-thyroxine). All participants were right-handed according to the Edinburgh Handedness Inventory (laterality index = 98.06, SD = 6.44) ^29^ and German native speakers. Prior to study inclusion, all participants gave written informed consent to the experimental procedure approved by the University of Tübingen local ethics committee. The study was conducted in accordance with the Declaration of Helsinki in its latest version.

### Adaptive 2-Back Paced Auditory Serial Addition Task (PASAT)

Participants were exposed to an adaptive, 2-back version of the Paced Auditory Serial Addition Task (PASAT) ^28^. Numbers ranging from 1 to 9 were continuously presented via headphones. Participants were asked to add the current number to the number presented before the previous one (2-back) and to type in their answer by pressing a correspondingly labeled keyboard button. Parallel to the next stimulus presentation, they received visual feedback, i.e. the screen flashed green for a correct answer and red for an incorrect, late or missed answer. The inter-stimulus interval between digit presentations adapted to participants’ performance. Initially set to 3 s, it was decreased by 0.1 s after four consecutive correct answers and increased 0.1 s after four consecutive wrong answers. The PASAT consisted of 16 practice trials followed by three task blocks lasting for 5 min, which were separated by breaks of 30 s. Due to the adaptive design the error percentage remained similar, although the number of correct trials could vary between task blocks.

### Positive and Negative Affect Schedule (PANAS)

To assess changes in negative affect during the experimental procedure, participants were administered the German version of the ‘Positive and Negative Affect Schedule’ (PANAS), a self-report to determine the participants’ current affective states ^30,31^. Ten positive and ten negative adjectives were rated on a five-point Likert scale ranging from 1 ‘not at all’ (in German: ‘gar nicht’) to 5 ‘very much’ (in German: ‘äußerst’). Participants completed the PANAS three times throughout each session: before starting the PASAT (pre), immediately after they completed the PASAT (post), and 90 min after task completion (follow up).

### Transcranial Direct Current Stimulation (tDCS)

A direct current of 1 mA was generated by a portable, battery-driven stimulator (NeuroConn GmbH, Illmenau, GER) and applied via a pair of 5 × 7 cm electrodes covered with conductive paste (Ten20^®^, Weaver and Company, Aurora, CO). The anodal electrode was placed over the left dorsolateral prefrontal cortex at F3 according to the international 10-20 system of electrode placement ^32^, whereas the cathodal reference electrode was fixated on the right upper arm over the deltoid muscle to prevent any opposite polarization of other brain regions ^33^. Two minutes before PASAT onset, the current was faded in for 5 s. During the anodal stimulation session, a continuous current of 1 mA was delivered for 20 min until task completion and then faded out for another 5 s. During sham stimulation, the current was only administered for 30 s before fading out. Impedance was controlled by the device and did not exceed 10 kΩ.

### Experimental Procedure

The experimental design was identical to Wiegand et al. (2019), where the behavioral data and changes in affect of the pilot cohort are described in more detail ^28^.

The study followed a single-blind, sham-controlled cross-over design. Each participant took part in two sessions with an interval of 7 days in between. To reduce variability, each session started at 2 PM. To ensure that the inclusion criteria were met, a brief screening including the Symptom-Checklist-90-Revised (SCL-90-R) to detect psychiatric symptoms and distress was performed in the first session ^34^. Apart from that, the two sessions only differed in the type of stimulation (anodal or sham) participants received. The order of stimulation was randomized and counterbalanced across participants.

Each session started with a saliva sampling. Then, a venous catheter was placed and the tDCS electrodes were fixated. Affective states were assessed (PANAS pre) and the instructions for the PASAT were given. The first blood sampling was done just before the stimulation started, but at least 15 min after the venous access had been established. Afterwards, participants were exposed to the 2-back PASAT while receiving tDCS (anodal or sham). Immediately after task completion, the second blood sample was collected and the PANAS (post) was administered. After removal of the electrodes, participants were exposed to relaxing music (genre: ambient electronic) via headphones until the end of the experiment, which was 90 min after task completion. Four more blood samples were collected 20, 40, 60 and 90 min after task completion. In addition, a second saliva sample was collected 30 min after the beginning of the 2-back PASAT. Finally, participants were administered the PANAS for a third time (follow-up). Supplementary Figure S5 depicts the experimental procedure graphically.

### DNAm Analysis

Blood samples were collected in EDTA tubes (2.7 ml Monovette^®^, Sarstedt, Sarstedt, GER), and stored at -80 °C. DNA was extracted with the QIAamp Blood Mini Kit (Qiagen, Hilden, GER) according to manufacturer’s instructions using 400 µl whole blood sample. To increase DNA yield, the final elution step was repeated using the 100 µl eluate of the first elution. DNA was quantified using the Qubit^®^ 2.0 Fluorometer (Life Technologies, Carlsbad, CA). Samples were stored at -20 °C.

500 ng of genomic DNA were bisulfite converted using the EpiTect Fast Bisulfite Kit (Qiagen, Hilden, GER) according to the manufacturer’s instructions. Bisulfite-converted DNA was eluted with 20 µl of the provided elution buffer. The purified bisulfite-converted DNA was stored at -20 °C.

A region-specific polymerase chain reaction (PCR) was performed for the *S-COMT* promoter region using the PyroMark PCR Kit (Qiagen, Hilden, GER) according to manufacturer’s instructions with previously published primers (F: 5-GAGTAGGTTGTGGATGGGTTGTA-3, R: 5-Biotin-ACATTTCTAAACCTTACCCCTCTA-3) ^35^. Successful amplification and specificity of the PCR products was verified and visualized via agarose gel electrophoresis.

DNAm was analyzed by pyrosequencing on a PyroMark Q24 system (Qiagen, Hilden, GER) using 5 µl biotinylated PCR product of each sample and a previously published sequencing primer (S: 5-GTAATATAGTTGTTAATAGTAGA-3) ^35^. DNAm level of the two investigated CpG sites (hg19 reference genome coordinates: chr22:19,950,054-19,950,064) was quantified using the PyroMark Q24 Software 2.0 (Qiagen, Hilden, GER). Each sample was analyzed twice and the mean percentage was used for further analysis. Samples with a deviation ≥ 3% between duplicates were repeated. To detect disparate amplification of unmethylated DNA fragments, a titration assay using standardized bisulfite-converted control DNA samples (EpiTect Control DNA, Qiagen, Hilden, GER) with established DNAm levels of 0%, 25%, 50%, 75% and 100% DNAm was performed.

### Saliva Cortisol Concentration

Saliva was sampled in Salivettes^®^ (Sarstedt AG & Co., Nümbrecht, GER) and stored at -80 °C. For analysis of cortisol levels, Salivettes^®^ were thawed and centrifuged for 2 min at 1000 g to collect saliva. Cortisol concentrations were determined using the Cortisol Saliva ELISA kit (IBL International, Hamburg, GER) according to manufacturer’s instructions. Cortisol concentrations were determined in duplicates and the mean coefficient of variation was below 10%.

### Statistical Analysis

All statistical calculations were performed using the software R (Version 3.5.1) ^36^ including the package nlme ^37^. The two cohorts were pooled for data analysis. Mean numbers of correct trials for each of the three task blocks were extracted from the adaptive 2-back PASAT as measure of task performance. For the PANAS questionnaire, mean scores were calculated for the 10 items comprising negative affect. DNAm levels of the *COMT* gene promoter region were expressed as the mean level of methylation of the two analyzed CpG sites.

Multilevel modeling was chosen over repeated-measures ANOVA to allow analyses of the effects of *stimulation* and *session* within the same statistical model. For all analyses (i.e., task performance, affect changes, DNAm changes and cortisol level changes) a multilevel model with the fixed effects *stimulation, session* and *time* (or *task block* accordingly in task performance data analyses) was estimated using maximum likelihood. A random intercept for each participant and random slopes for the effects of *session* and *time* (or *task block*) were included to account for individual differences in the outcome variable, in the effect of *session* and in the effect of *time* (or *task block*) within each session. The error term was modeled as a first order autoregressive process to account for serial autocorrelations due to the repeated measures design. The severity of multicollinearity was assessed by the variance inflation factor (VIF). To assure a VIF < 10, the interaction of *stimulation* and *session* and, hence, the three-way interaction of *stimulation, session* and *time* (or *task block*) was eliminated from the models for task performance, DNAm and cortisol levels ^38,39^. For changes in affect, a full model was estimated as VIF < 10 was given for all predictors. A linear model was fitted for the analyses of task performance, DNAm and cortisol levels, whereas a quadratic term for time was included in the model for affect resulting in a better fit for changes over time. Unstandardized (B) as well as standardized (β) parameter estimates were reported and statistical significance was assessed at *p* < 0.05. Additionally, an analysis of variance table for each model is given in the Supplementary Tables S1 – S4 reporting the overall significance of all terms ^40^. Post-hoc pairwise comparisons were performed after significant effects.

Similarly, a multilevel model was fitted with the fixed effects *stimulation in first session* and *session* including only DNAm data of the first time point of each session, to examine whether DNAm changes induced in the first session were preserved until the second session with respect to the type of stimulation received during the first session. Random intercepts estimated for each participant and random slopes for the effect of session were included to account for individual variance.

Finally, Pearson’s correlation was used to test for an interrelation between changes in DNAm levels (*COMT*-methylation_post90_ - *COMT*-methylation_pre_) and cortisol concentration (cortisol_post_ – cortisol_pre_) and between changes in negative affect (negative affect_post_ – negative affect_pre_) and in cortisol concentration during the first session.

## Results

### Study Sample

Participants were randomly assigned to the order of stimulation (anodal/sham or sham/anodal) they received during the two experimental sessions. The two resulting groups showed no significant differences with respect to age (*t*(40)=1.52, *p*=0.14), years of formal education (*t*(40)=1.06, *p*=0.30), math performance at school (*t*(36)=-0.54, *p*=0.59), body mass index (*t*(40)=-0.24, *p*=0.81), global severity index (SCL-90-R) (*t*(40)=0.59, *p*=0.56), or *COMT* Val108/158Met genotype (χ^2^=0.15, *p*=0.93). A more detailed description of the sample characteristics including sociodemographic variables and information on the *COMT* Val108/158Met genotype can be found in Supplementary Table S5.

### Task Performance

Task performance in the adapted version of the PASAT was evaluated by a linear mixed model with the predictors *stimulation* (anodal, sham), *session* (1, 2) and *task block* (1, 2, 3). *Task block* (B=3.80, SE=0.99, β=0.24, *t*(202)=3.84, *p*<0.001) and *session* (B=9.53, SE=0.79, β=0.60, *t*(202)=12.07, *p*<0.001) significantly predicted task performance due to an increasing number of correct trials over the course of the three task blocks within each session and from the first to the second session, respectively. Furthermore, the interaction of *session* and *task block* predicted task performance significantly (B=-2.36, SE=1.12, β=0.15, *t*(202)=-2.11, *p*=0.036). Neither *stimulation* (B=-0.39, SE=0.97, β=-0.02, *t*(202)=-0.40, *p*=0.69), nor the interaction between *stimulation* and *task block* (B=1.54, SE=1.29, β=0.10, *t*(202)=1.19, *p*=0.23) predicted task performance significantly.

Follow-up t-tests showed significant increases in the number of correct trials from task block 1 to task block 2 (*t*(41)=-2.86, *p*=0.007, |d|=0.44), from task block 2 to task block 3 (*t*(41)=- 3.20, *p*=0.003, |d|=0.49) and from task block 1 to task block 3 (*t*(41)=-5.94, *p*<0.001, |d|=0.92) during session 1. During session 2 there was a significant increase from task block 1 to task block 3 (*t*(41)=-2.63 *p*=0.012, |d|=0.41) and from task block 2 to task block 3 (*t*(41)=-2.98 *p*=0.005, |d|=0.46), but not from task block 1 to task block 2 (*t*(41)=-0.33 *p*=0.74). Figure 1 depicts the number of correct trials for each task block during each session with regard to stimulation condition.

**Figure 1.**
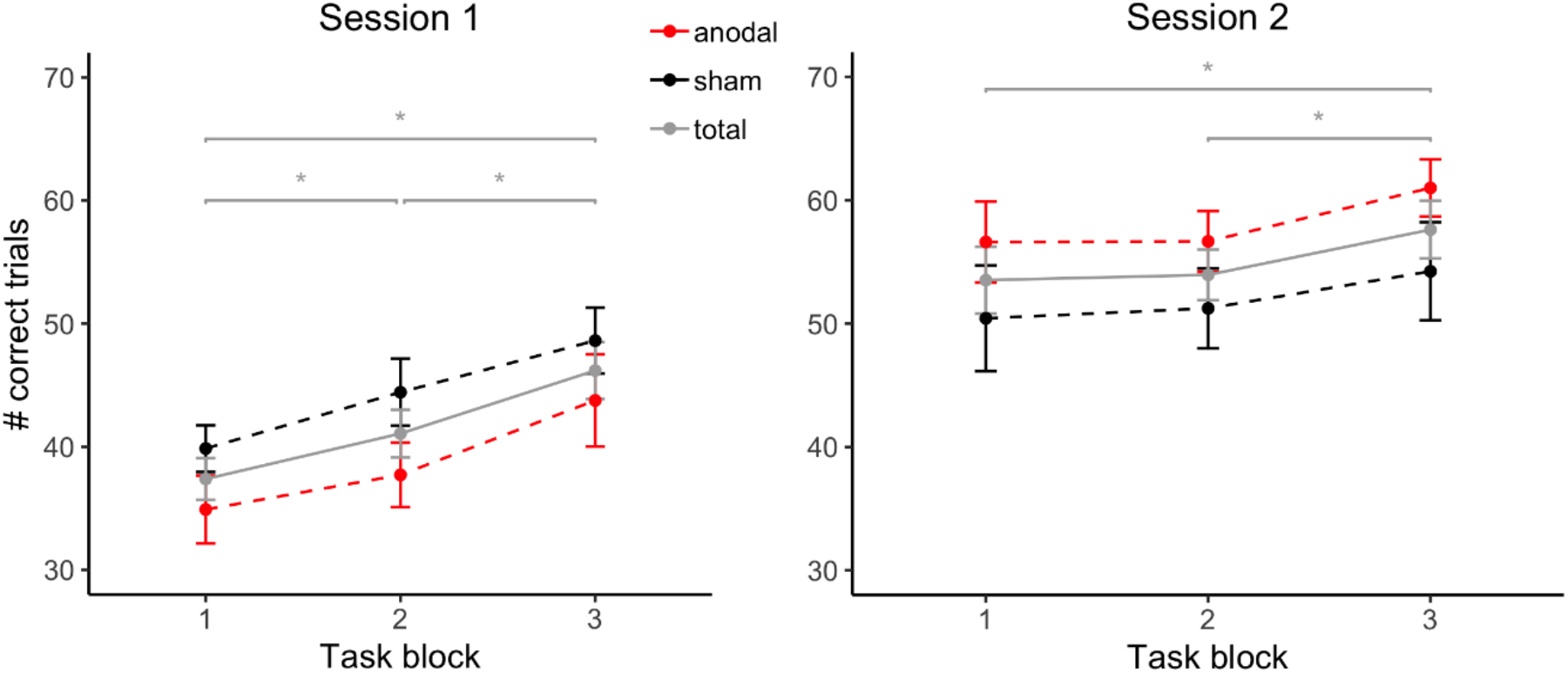
Task performance during each session with regard to stimulation condition. As the order of received stimulation (‘anodal - sham’ or ‘sham - anodal’) was a between-subject factor, participants receiving anodal stimulation during the first session (n = 21) received sham stimulation during their second session and vice versa (n = 21). Error bars depict standard errors of the mean.

### Affective Changes

Changes in negative affect were investigated by a multilevel mixed model with the predictors *stimulation* (anodal, sham), *session* (1, 2) and *time* (pre, post, follow-up).

*Session* (B=-0.09, SE=0.04, β=-0.38, *t*(199)=-2.45, *p*=0.015) and *time* (B=-0.12, SE=0.03, β=- 0.53, *t*(199)=-3.84, *p*<0.001) both significantly predicted changes in negative affect. While the interaction of *stimulation* and *time* did not predict negative affect significantly ((B=0.07, SE=0.04, β=0.29, *t*(199)=1.59, *p*=0.11), the interaction of *session* and *time* predicted the outcome variable significantly (B=0.14, SE=0.05, β=0.60, *t*(199)=3.05, *p*=0.003). Furthermore, the three-way interaction of *stimulation, session* and *time* predicted negative affect by trend (B=-0.12, SE=0.07, β=-0.54, *t*(199)=-1.85, *p*=0.065), indicating that the effect of tDCS on changes in negative affect might be different in the two sessions.

In the first session, participants receiving sham stimulation showed an increase in negative affect by trend (*t*(20)=-1.95, *p*=0.066, |d|=0.42), whereas the negative affect did not change in participants under anodal stimulation (*t*(20)=-0.38, *p*=0.71). There were no changes in negative affect during the second session, neither for participants under anodal stimulation (*t*(20)=-1.16, *p*=0.26), nor for participants in the sham condition (*t*(20)=0.13, *p*=0.90). Figure 2 depicts the changes in negative affect during each session with regard to stimulation condition.

**Figure 2.**
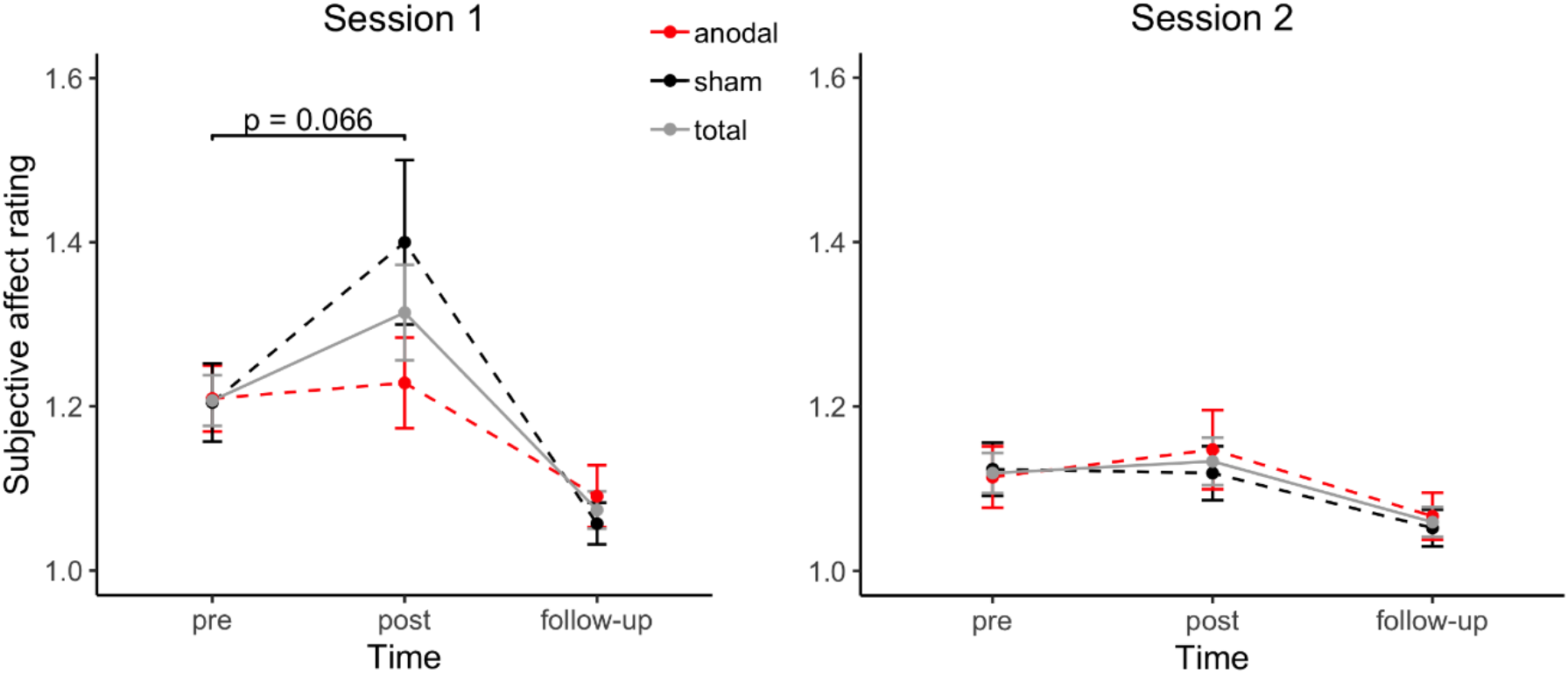
Changes in negative affect during each session with regard to stimulation condition. Subjective rating of negative affect is shown separately for each session in pre- and post-task and follow-up condition. As the order of received stimulation (‘anodal - sham’ or ‘sham - anodal’) was a between-subject factor, participants receiving anodal stimulation during the first session (n = 21) received sham stimulation during their second session and vice versa (n = 21). Error bars depict standard errors of the mean.

### DNAm Changes in *COMT* Gene Promoter Region

In a linear mixed model with the predictors *stimulation* (anodal, sham), *session* (1, 2) and *time* (hours), *stimulation* (B=0.98, SE=0.42, β=0.13, *t*(457)=2.34, *p*=0.020) and *time* (B=0.65, SE=0.17, β=0.06, *t*(457)=3.75, *p*<0.001) predicted DNAm levels significantly, implying an effect of tDCS on the DNAm and dynamic DNAm changes during the experimental procedure. While the interaction of *stimulation* and *time* (B=-0.09, SE=0.23, β=-0.01, *t*(457)=-0.37, *p*=0.71) did not predict the outcome variable significantly, the interaction of *session* and *time* significantly predicted DNAm levels (B=-0.64, SE=0.17, β=-0.05, *t*(457)=-3.77, *p*<0.001). This indicates that the effect of the PASAT performance on *COMT* DNAm over time differs between the first and the second session and that the tDCS effect occurs between and not within the interventions.

Follow-up t-tests showed a significant increase in DNAm during session 1 from time point pre (57.70% methylated) to post90 (59.33% methylated) disregarding the stimulation condition (*t*(41)=-4.30, *p*<0.001, |d|=0.66), driven by an almost continuous increase in DNAm levels during the experimental procedure. Further t-tests comparing ‘pre’ with all post time points during the first session, showed that this increase is significant from time point post20 (58.69% methylated) onwards (*t*(41)<-2.48, *p*<0.017, |d|>0.38). For session 2, no change in DNAm from time point pre to any post time point was observed (|*t*(41)|<1.8, *p*>0.08). Figure 3A depicts changes in DNAm over the experimental procedure separately for the two sessions.

**Figure 3.**
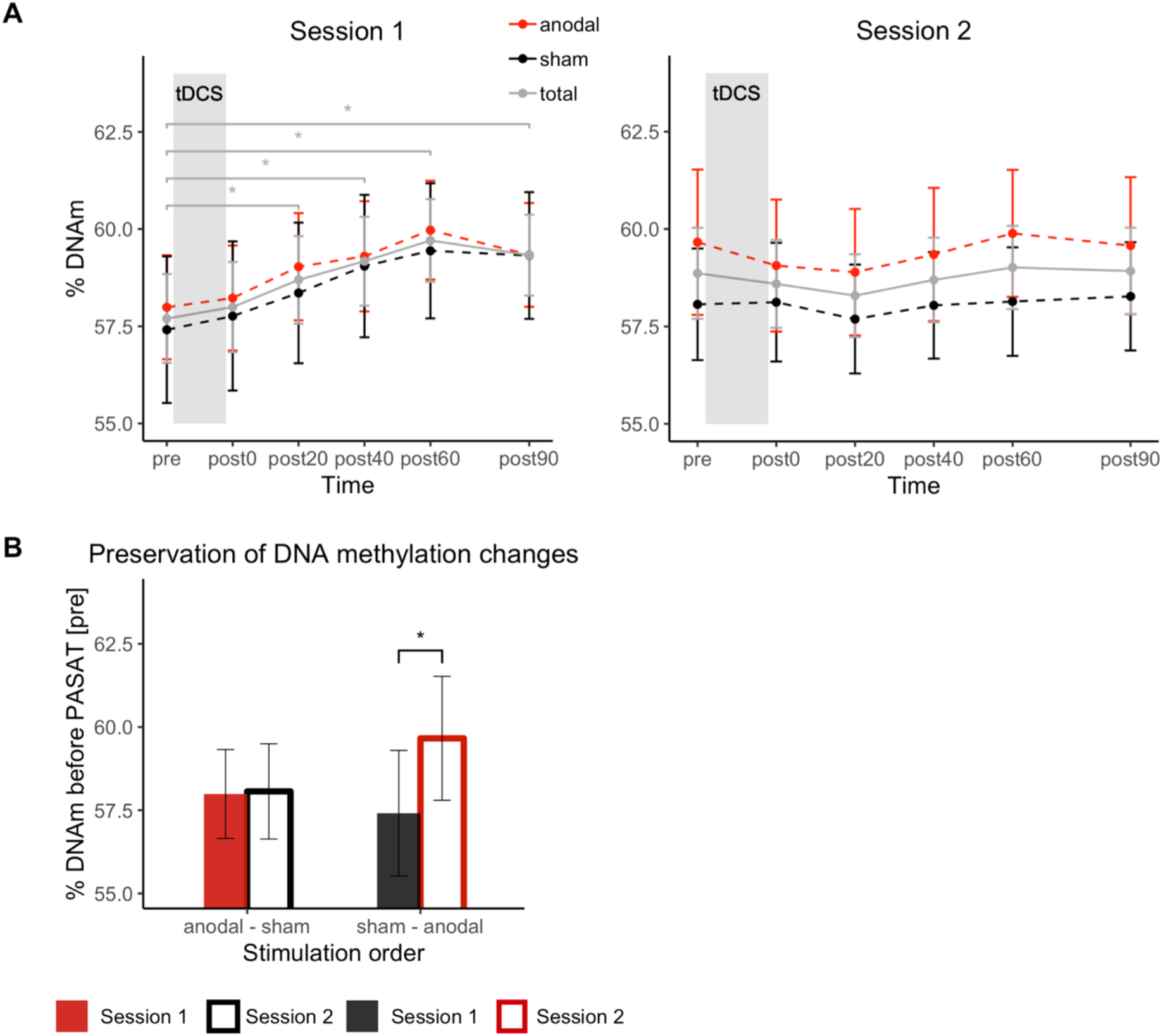
DNAm changes during each session with regard to stimulation condition and its preservation over one week. A. % DNAm is shown separately for the six time points during each session. Each participant in the anodal stimulation group in session 1 (n = 21) was receiving sham stimulation in session 2 and vice versa (n = 21). B. % DNAm for session 1 and 2 at time point ‘pre’ grouped by order of stimulation conditions (‘anodal - sham’ (n = 21) or ‘sham - anodal’ (n = 21)). The figure illustrates the comparison of % DNAm before (‘pre’) the first (session 1) and second (session 2) PASAT training within subjects who received tDCS (‘anodal - sham’) and subjects who did not receive effective tDCS in session 1 (‘sham - anodal’). Error bars depict standard errors of the mean.

Demonstrating that the increase in DNAm during session 1 was still present in session 2, i.e. that it was preserved over 1 week, a multilevel model was fitted with the predictors *stimulation in first session* (anodal, sham) and *session* (1, 2) including only DNAm data of the first time point of each session. The interaction of *stimulation in first session* and *session* significantly predicted DNAm levels at the beginning of each session (B=1.54, SE=0.68, β=0.21, *t*(40)=2.26, *p*=0.030). As depicted in Figure 3B, the increase in DNAm during the first session could still be detected in the beginning of session 2 in sham-treated participants (*t*(20)=-2.95, *p*=0.008, |d|=0.64), but not in anodal-treated participants (*t*(20)=-0.13, *p*=0.90).

### Cortisol Changes

In a linear mixed model with the predictors *stimulation* (anodal, sham), *session* (1, 2) and *time* (pre, post), *session* (B=-0.08, SE=0.02, β=-0.28, *t*(121)=-3.40, *p*<0.001) predicted cortisol levels significantly and *time* predicted cortisol levels by trend (B=0.05, SE=0.03, β=0.18, *t*(121)=1.90, *p*=0.060). Neither *stimulation* (B=0.02, SE=0.03, β=0.06, *t*(121)=0.76, *p*=0.45) nor the interaction of *stimulation* and *time* (B=-0.02, SE=0.03, β=-0.06, *t*(121)=-0.54, *p*=0.59) significantly predicted the outcome variable. However, the interaction of *session* and *time* predicted cortisol levels significantly (B=-0.07, SE=0.03, β=-0.25, *t*(121)=-2.60, *p*=0.011), indicating differences in cortisol changes in the two sessions.

Follow-up t-tests showed a significant increase in cortisol levels during session 1 (*t*(41)=-2.50, *p*=0.017, |d|=0.39), while no cortisol changes were detected in session 2 (*t*(41)=-0.26, *p*=0.80). Figure 4 depicts changes in cortisol levels during each session.

**Figure 4.**
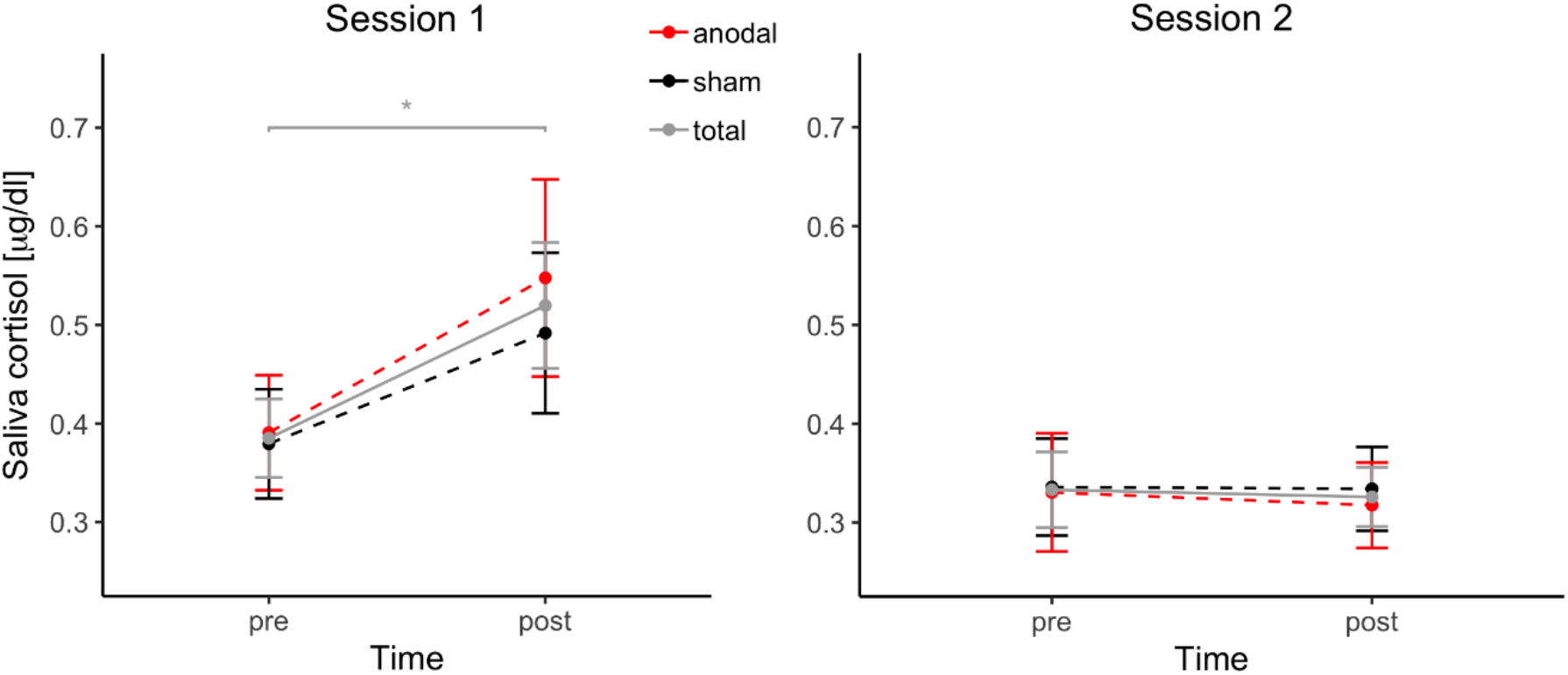
Cortisol concentration changes during each session with regard to stimulation condition. Saliva cortisol levels are shown separately for each session in pre- and post-task condition. As the order of received stimulation (‘anodal - sham’ or ‘sham - anodal’) was a between-subject factor, participants receiving anodal stimulation during the first session (n = 21) received sham stimulation during their second session and vice versa (n = 21). Error bars depict standard errors of the mean.

### Correlation of DNAm Changes and Cortisol Changes

During the first session, there was a significant correlation between changes in cortisol concentration and changes in DNAm levels (r=0.359, *p*=0.019) as depicted in Figure 5. There was no correlation between changes in cortisol concentration and changes in negative affect (r=-0.139, p=0.38).

**Figure 5.**
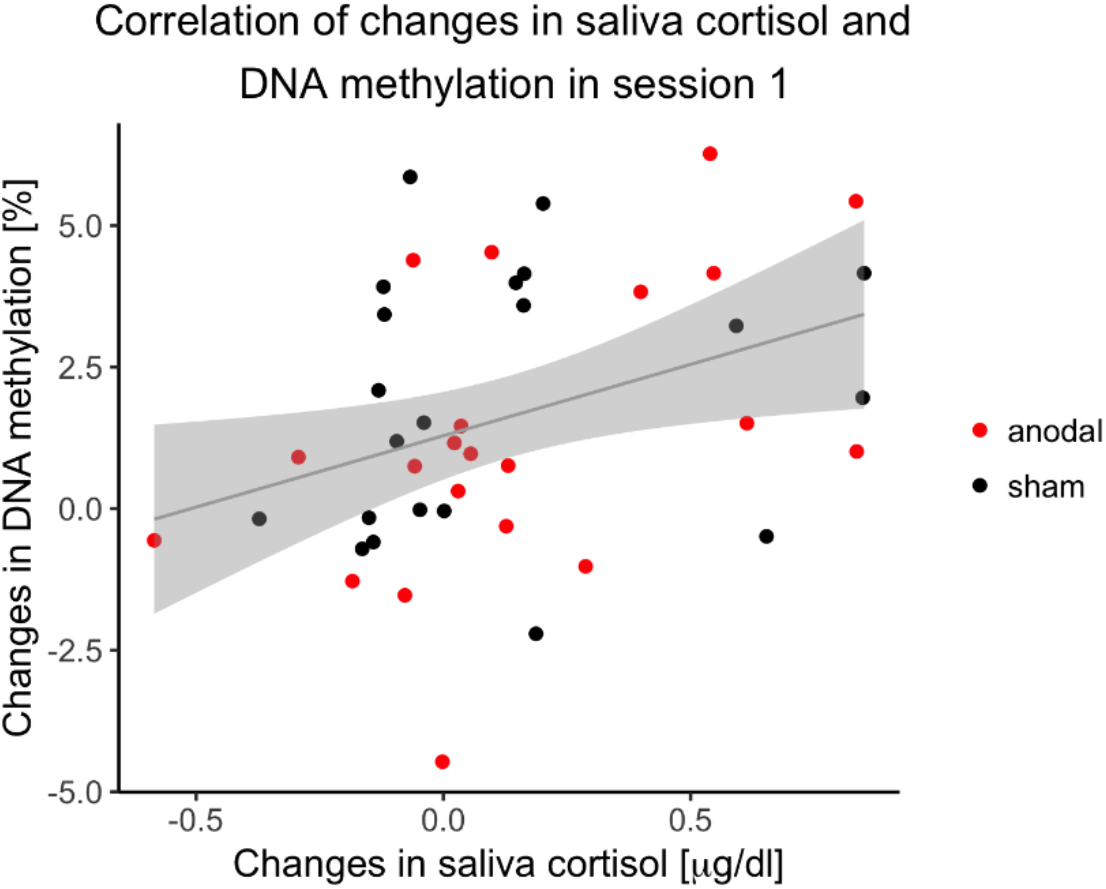
Correlation of DNAm changes and cortisol changes. Correlation of changes in DNAm during session 1 with changes in saliva cortisol concentration (n = 42). Regression line with 0.95 confidence interval.

## Discussion

The key findings of the present study are i) a continuous increase of *COMT* gene promotor methylation in blood after a challenging, stressful and frustrating cognitive task which correlates with an increase in salivary cortisol, ii) increased *COMT* DNAm detectable one week later, and iii) a suppression of this lasting effect by concurrent activity-enhancing anodal tDCS to the dorsolateral prefrontal cortex. These data support the notion of dynamic DNAm in response to mental stress that is associated with changes in cortisol levels and can be modulated by tDCS.

To date, few studies have reported dynamic changes in DNAm in response to a mental stress paradigm. In fact, DNAm levels have been regarded as rather stable, long-term epigenetic marks in somatic cells, which might even be maintained over numerous cell divisions ^41^. However, previous studies have associated differential DNAm patterns with diseases like post-traumatic stress disorder, which can be triggered by the experience of a single stressful life event ^42^. Yet, little is known about the time frame of the formation of these methylation changes and the amount of stress load required to induce these changes. An increase in *OXTR* DNAm was observed already 10 min after the exposure to the TSST ^11^ and changes in *FKBP5* gene expression within 70 min ^43^. Congruent with these studies, our data add evidence to those immediate effects on DNAm and, moreover, the continuous increase in *COMT* promoter methylation over the course of five independent measurements within 90 minutes after stress exposure makes a random variation rather unlikely.

The correlation of changes in saliva cortisol and DNAm supports the idea that the stress hormone cortisol links stress with DNAm changes detectable in peripheral blood. This is also in line with previous findings reporting DNAm differences after glucocorticoid exposure ^44,45^. We specifically chose to investigate methylation dynamics of the *COMT* gene, as several studies indicate its role in stress reactivity on a genetic ^46,47^ and epigenetic level ^48^. Furthermore, the *COMT* gene interacts with cognitive performance and tDCS effects, supporting its eligibility as candidate gene ^16,49^. Due to the inaccessibility of living brain tissue we investigated DNAm differences in whole blood. This leads to the question to what extent blood DNAm can serve as proxy marker for DNAm in neuronal tissue. Indeed, for *COMT* DNAm several studies show that methylation status in peripheral tissue can serve as surrogate for brain DNAm ^48,50^. More importantly, our data show that changes in DNAm correlate with changes in cortisol concentration. Saliva cortisol correlates well with the concentration of free circulating cortisol ^51^. Therefore, it is possible that the dynamic DNAm is mediated by alterations in plasma cortisol levels. Since cortisol can cross the blood brain barrier, it might elicit similar effects on DNAm in neuronal tissue. Taken together, these data support the hypothesis that dynamic epigenetic modifications are invoked immediately by exposure to stress and associated with the released cortisol.

The COMT enzyme is involved in the degradation of catecholamines like dopamine and changes in its expression levels could consequently affect catecholamine concentrations. There are two isoforms of COMT regulated by two different promotors. The predominant form in peripheral tissue is the soluble isoform (S-COMT) ^52^. The CpG sites investigated in our study are located within its promotor region and, hence, may be involved in the regulation of *S-COMT* expression. However, since the observed DNAm differences were relatively small, their functional relevance needs to be clarified in future studies including gene expression data. Furthermore, in the brain the primarily expressed mRNA is encoding the membrane-bound isoform (MB-COMT), however, there is evidence that, to a lower extent, the S-COMT form is also expressed ^52^. The investigated CpG sites fall within the gene body region of *MB-COMT*. Therefore, if *COMT* DNAm is affected to a similar extent in neuronal tissue, effects on *COMT* expression levels in the brain might be diverse ^53^. Given this limitation, conclusions about a potential epigenetic feedback loop controlling prefrontal dopamine activity as well as neuroplasticity and behavior is beyond the scope of this study. The dynamics of stress-related epigenetic changes and its relation to the neurotransmitter metabolism in the brain are likely better suited to animal studies.

Considering that the PASAT only presents a relatively mild stressor, it is quite remarkable that it elicited significant changes in DNAm which even persisted over one week. Of note, the PASAT task is originally designed as a measure of information processing ability ^54^. However, previous studies have also demonstrated that the PASAT induces psychological stress ^55^. In this study, a more challenging design (2-back task) was used, probably leading to an even more stressful experience while performing this not only cognitively but also emotionally challenging task. This is supported by the increase in cortisol levels after exposure to the first task. Interestingly, no changes in cortisol levels were observed during the second session. Probably, participants are adapting to the task, which is why a stress response is only elicited when the task is unfamiliar. Being already mentally prepared to encounter a difficult task that comes along with frequent negative feedback might attenuate the stress experience. Although this indicates the limited validity of the PASAT as a mere stress task when used repeatedly, it circumvents the necessity of a less stressful control task. It controls already for the possibility that any other parameter, like, for example, the physical stress of the venous catheter placement, might have led to the observed changes.

Similar to the observed changes in cortisol and DNAm levels, there was also an increase in negative affect by trend during the first, but not during the second experimental session. However, in contrast to the molecular markers, these changes in affect seemed to be suppressed by anodal tDCS. As this short-term effect of tDCS on the affective experience is not observed in DNAm and cortisol levels, a correlation between changes in affect and cortisol is missing. The at least trend-wise effect of tDCS on negative affect is in line with the hypothesis that activity enhancing stimulation applied over the left dorsolateral prefrontal cortex increases cognitive control over emotion and, thereby, leads to stabilization in affect ^56,57^.

Furthermore, the application of tDCS during the first session affected the preservation of DNAm changes. While in sham-treated participants an increase in DNAm was still detectable one week later, methylation levels returned to baseline when anodal stimulation had been applied. Although we were not able to detect a significant difference between the active and sham stimulated subjects in salivary cortisol levels 30 min after intervention, it is possible that these differed in the cumulated amount of cortisol excretion over the time of the experiment. This was not captured by our experimental design, but a modulation of the stress response by brain stimulation has been reported previously ^58,59^, and may have been involved in mediating the observed differences in the stability of stress-induced DNAm changes. It has been previously shown in a mouse model that activity enhancing tDCS can lead to alterations in histone modifications, chromatin remodeling and changes in gene expression ^60^. Our data provide preliminary evidence for a lasting modulation of DNAm changes by tDCS suggesting that tDCS induces system-wide effects detectable in peripheral tissues like whole blood through intermediary pathways.

One confounding factor which might affect DNAm are alterations in blood cell composition. Since epigenetic patterns are cell type specific, DNAm measured in whole blood might be influenced by cell type composition. According to the iMETHYL database, which provides cell type specific DNAm patterns based on a Japanese population, the methylation level of the investigated CpG sites in the *COMT* gene promoter region is very similar between neutrophils and monocytes but substantially lower for CD4-positve T-lymphocytes ^61-63^. Studies investigating stress-induced immunological reactions report changes in leukocyte counts after severe physical or acute psychological stress which is often accompanied by increased glucocorticoid levels ^64^. Whether these effects can already occur within a narrow time frame of two hours is disputed and potentially species specific ^65^. For humans, there is evidence that cortisol induced effects on leukocyte composition occur with a time lag of two hours ^66,67^. Therefore, we believe that it is rather unlikely that the increase in *COMT* promotor methylation already present 20 min after stress exposure reported here is merely a secondary effect of changes in cell type composition. Nevertheless, controlling for potential cell type composition effects in future studies will be reasonable.

The novel findings of the present study evoke several lines of research to validate and extend our results. Previous research suggests that activity-enhancing stimulation applied over the left dorsolateral prefrontal cortex increases cognitive control over emotion and, thereby, leads to stabilization in affect ^56,57^. Since task-induced changes in negative affect missed significance in our study, future studies using well-established mental stress paradigms with greater power to induce changes in affect are needed to investigate the relation between tDCS effects on emotion regulation, cortisol concentration changes, and the persistence of DNAm changes. Furthermore, longitudinal studies with several sessions of tDCS application in the context of mental stress may help to investigate long-term effects of tDCS treatment on DNAm. Previous studies have shown that DNAm might not only change in association with stress exposure but also in response to different psychological treatments such as cognitive-behavioral therapy (CBT) ^68^. Given that our data suggest that tDCS reduces potentially maladaptive epigenetic effects in response to stressful experiences, it might be a valuable addition to complement conventional therapies such as exposure-based CBT.

Another limitation of our study concerns the sample size. For tDCS effects, we included an adequate number of participants as we can assume small to intermediate effect sizes ^69^. However, since this is the first human study investigating the effects of tDCS on DNAm changes, it was an explorative approach to detect potential effects and provide a first estimation of effect sizes for future research.

In the field of molecular psychiatry, the inaccessibility of living brain tissue is a major challenge. However, our findings support the notion of DNAm status in whole blood as a potential biomarker in the context of stress-related behavior, most likely via intermediary hormones. On that account, it is reasonable to study peripheral tissue in order to track epigenetic traces of a stressful cognitive effort in behaving human subjects, with due caution regarding the direct transferability to changes in the brain. Finally, by showing that prefrontal tDCS can affect the stability of stress-induced DNAm changes in a gene regulating neurotransmission, these data point towards possible alternative pathways that might be involved in the therapeutic effects of brain stimulation.

## Declarations

### Ethics Approval and Consent to Participate

The study was performed in accordance with the Declaration of Helsinki and ethical approval was obtained at the University of Tübingen local ethics committee. All participants gave written informed consent to participate in the study.

### Availability of Data and Materials

The datasets generated and analyzed during the current study are available from the corresponding author on reasonable request.

### Competing Interest

The authors declare no conflict of interest.

### Funding

This work was supported by the GCBS research consortium (FKZ 01EE1403D) funded by the Federal Ministry of Education and Research.

### Authors’ Contributions

A.W., V.N. and C.P. designed the experiments; A.W. and A.B. ran the experiments; A.W. and S.W. analyzed the data; A.W., V.N. and C.P. wrote the manuscript; C.B., S.W. and A.B. commented on the manuscript.

## Acknowledgements

We acknowledge support by the Federal Ministry of Education and Research and Open Access Publishing Fund of University of Tübingen. We thank Dr. Daniel Bucher for proofreading the manuscript. The authors also wish to express their appreciation to all participants.

## Notes

### Competing Interest Statement

The authors have declared no competing interest.

