## Supplementary for "Dynamic DNA methylation changes in the *COMT* gene promoter region in response to mental stress and its modulation by transcranial direct current stimulation"

### Supplementary Material

#### Data from pilot cohort only (n = 22)

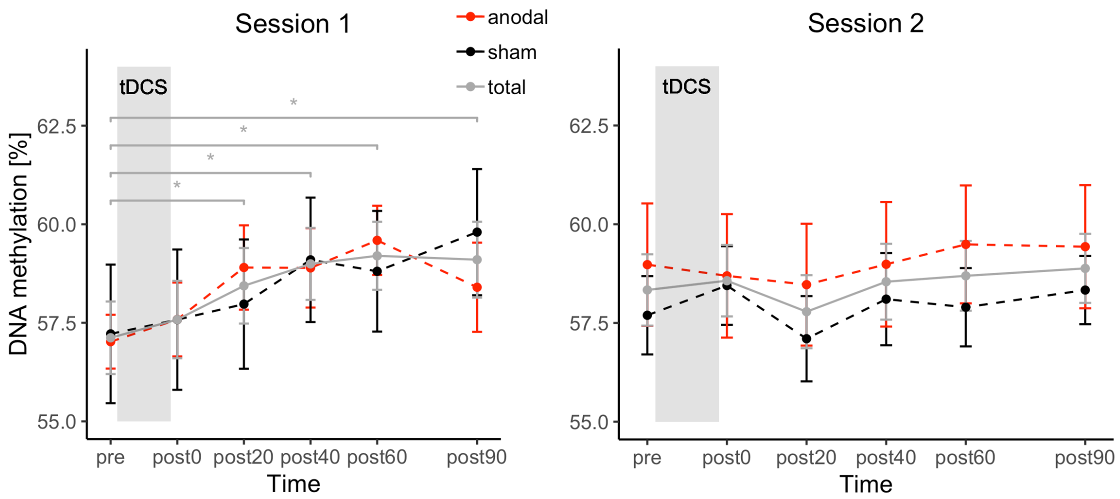


**Supplementary Figure S1. DNA methylation changes in pilot cohort during each session with regard to stimulation condition.** % DNA methylation is shown separately for the six time points during each session. Each participant in the anodal stimulation group in session 1 (n = 11) was receiving sham stimulation in session 2 and vice versa (n = 11). Error bars depict standard errors of the mean.

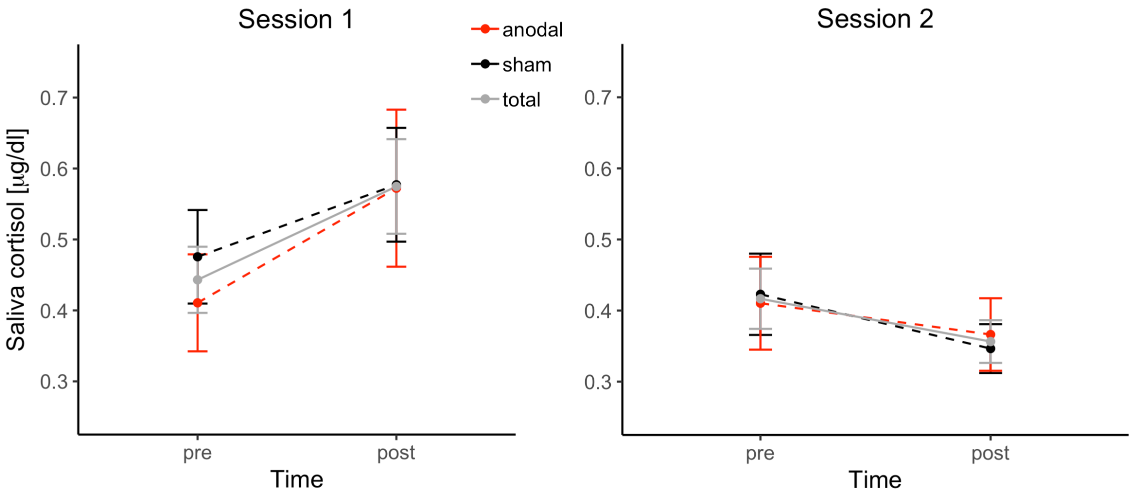


**Supplementary Figure S2. Cortisol concentration changes in pilot cohort during each session with regard to stimulation condition.** Saliva cortisol levels are shown separately for each session in pre- and post-task condition. As the order of received stimulation (anodal/sham or sham/anodal) was a between-subject factor, participants receiving anodal stimulation during the first session (n = 11) received sham stimulation during their second session and vice versa (n = 11). Error bars depict standard errors of the mean.

#### Data from replication cohort only (n = 20)

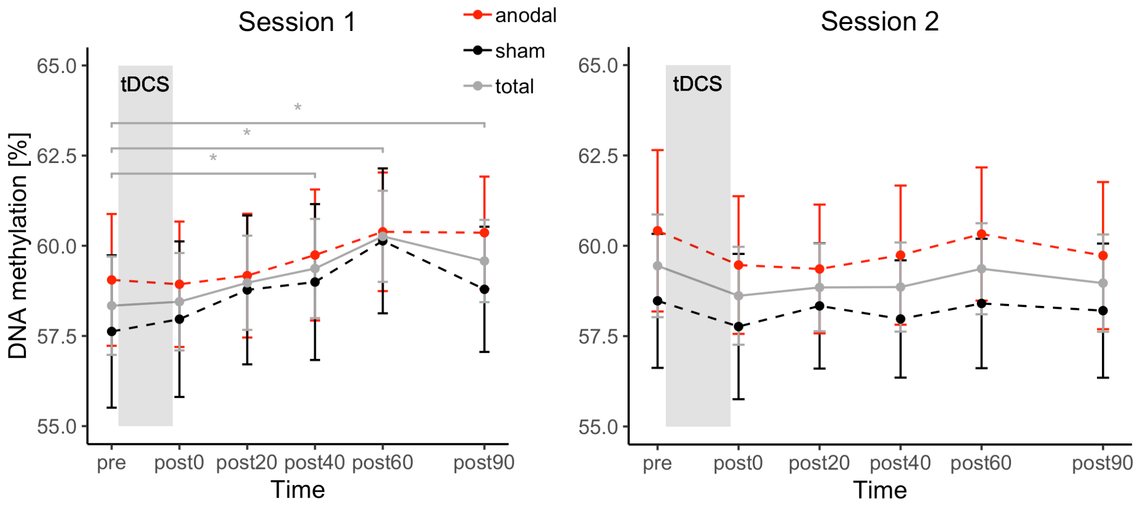


**Supplementary Figure S3. DNA methylation changes in replication cohort during each session with regard to stimulation condition.** % DNA methylation is shown separately for the six time points during each session. Each participant in the anodal stimulation group in session 1 (n = 10) was receiving sham stimulation in session 2 and vice versa (n = 10). Error bars depict standard errors of the mean.

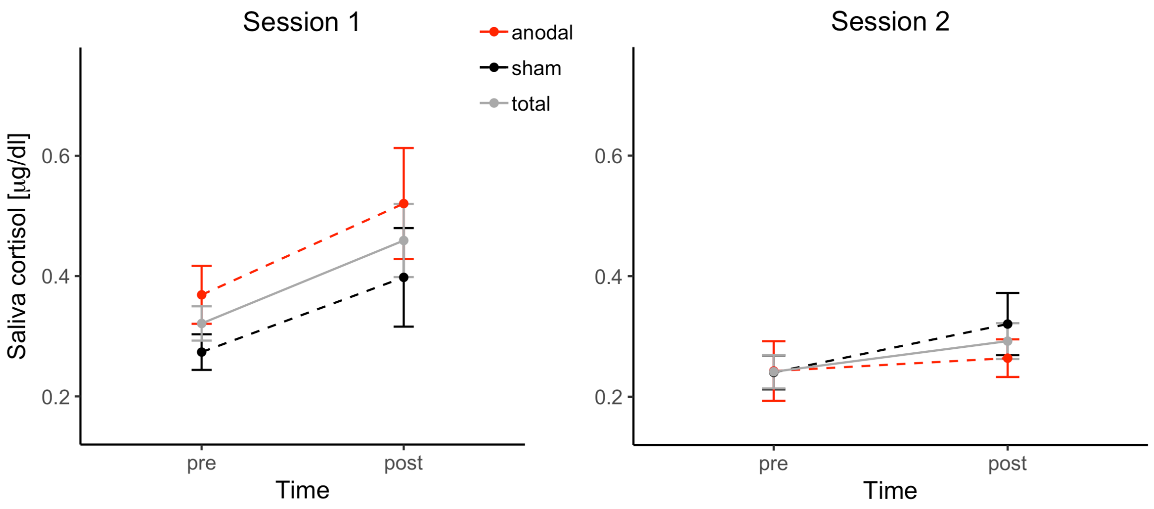


**Supplementary Figure S4. Cortisol concentration changes in replication cohort during each session with regard to stimulation condition.** Saliva cortisol levels are shown separately for each session in pre- and post-task condition. As the order of received stimulation (anodal/sham or sham/anodal) was a between-subject factor, participants receiving anodal stimulation during the first session (n = 10) received sham stimulation during their second session and vice versa (n = 10). Error bars depict standard errors of the mean.

### Experimental Procedure

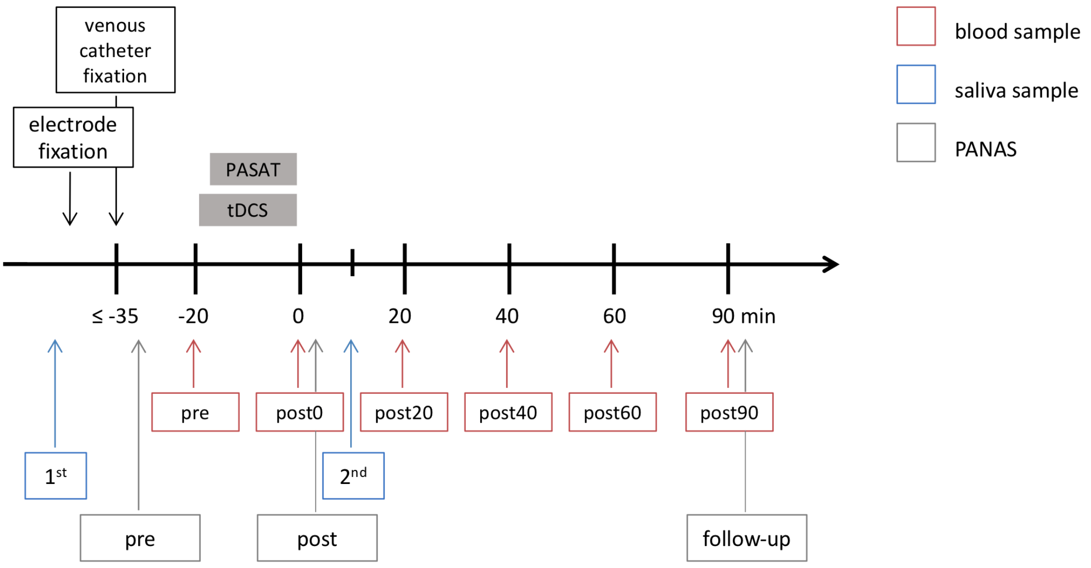


**Supplementary Figure S5. Experimental procedure.** Schematic representation of an experimental session.

### Analysis of Variance Tables

**Supplementary Table S1. Task performance**

| Term | *F* | *p* |
|---|---|---|
| **Intercept** | $F(1,202) = 776.10$ | $p < 0.001$ |
| **Stimulation** | $F(1,202) = 0.17$ | $p = 0.68$ |
| **Session** | $F(1,202) = 142.71$ | $p < 0.001$ |
| **Task block** | $F(2,202) = 5.40$ | $p = 0.005$ |
| **Stimulation * task block** | $F(2,202) = 0.83$ | $p = 0.44$ |
| **Session * task block** | $F(2,202) = 2.33$ | $p = 0.10$ |

**Supplementary Table S2. Affect changes**

| Term | *F* | *p* |
|---|---|---|
| **Intercept** | $F(1,199) = 5366.20$ | $p < 0.001$ |
| **Stimulation** | $F(1,199) = 0.11$ | $p = 0.75$ |
| **Session** | $F(1,199) = 2.75$ | $p = 0.09$ |
| **Time** | $F(2,202) = 11.02$ | $p < 0.001$ |
| **Stimulation * session** | $F(1,199) = 0.28$ | $p = 0.60$ |
| **Stimulation * time** | $F(2,202) = 1.17$ | $p = 0.31$ |
| **Session * time** | $F(2,199) = 7.18$ | $p = 0.001$ |
| **Stimulation * session * time** | $F(2,199) = 1.72$ | $p = 0.18$ |

**Supplementary Table S3. DNA methylation changes**

| Term | *F* | *p* |
|---|---|---|
| **Intercept** | $F(1,457) = 3876.71$ | $p < 0.001$ |
| **Stimulation** | $F(1,457) = 5.89$ | $p = 0.016$ |
| **Session** | $F(1,457) = 0.41$ | $p = 0.52$ |
| **Time** | $F(1,457) = 17.09$ | $p < 0.001$ |
| **Stimulation * time** | $F(1,457) = 0.14$ | $p = 0.71$ |
| **Session * time** | $F(1,457) = 14.25$ | $p < 0.001$ |

**Supplementary Table S4. Cortisol concentration changes**

| Term | *F* | *p* |
|---|---|---|
| **Intercept** | *F(1,121)* = 6.33 | *p* = 0.013 |
| **Stimulation** | *F(1,121)* = 2.51 | *p* = 0.12 |
| **Session** | *F(1,121)* = 6.12 | *p* = 0.014 |
| **Time** | *F(1,121)* = 0.38 | *p* = 0.54 |
| **Stimulation * time** | *F(1,121)* = 0.01 | *p* = 0.93 |
| **Session * time** | *F(1,121)* = 6.75 | *p* = 0.011 |

### Sample description

**Supplementary Table S5.** Sample characteristics

| | **Group sham – anodal (n=21)** | **Group anodal – sham (n=21)** |
|---|---|---|
| **Age [years]** | 22.71 (± 2.81; 18-28) | 24.19 (± 3.44; 18-29) |
| **Years of education [years]** | 16.38 (± 2.99; 12-22) | 17.40 (± 3.29; 12-23) |
| **Math performance at school*** | 10.76 (± 2.47; 5-15) | 10.23 (± 2.93; 5-15) |
| **Laterality index**** | 99.05 (± 4.37; 80-100) | 97.08 (± 8.00; 70-100) |
| **BMI [kg/m$^2$]** | 23.00 (± 2.10; 19.59-27.44) | 22.86 (± 1.83; 19.94-25.98) |
| **SCL-90-R** | | |
| Somatization | 0.16 (± 0.18; 0.00-0.58) | 0.21 (± 0.22; 0.00-0.75) |
| Obsessive-compulsive | 0.41 (± 0.36; 0.00-1.20) | 0.38 (± 0.29; 0.00-1.00) |
| Interpersonal sensitivity | 0.18 (± 0.19; 0.00-0.67) | 0.22 (± 0.28; 0.00-1.00) |
| Depression | 0.19 (± 0.22; 0.00-0.69) | 0.27 (± 0.27; 0.00-1.00) |
| Anxiety | 0.18 (± 0.13; 0.00-0.40) | 0.11 (± 0.13; 0.00-0.40) |
| Anger-hostility | 0.19 (± 0.22; 0.00-0.83) | 0.21 (± 0.26; 0.00-0.83) |
| Phobic anxiety | 0.05 (± 0.12; 0.00-0.43) | 0.01 (± 0.03; 0.00-0.14) |
| Paranoid ideation | 0.12 (± 0.21; 0.00-0.83) | 0.20 (± 0.33; 0.00-1.17) |
| Psychoticism | 0.06 (± 0.09; 0.00-0.30) | 0.15 (± 0.21; 0.00-0.70) |
| GSI | 0.18 (± 0.12; 0.00-0.44) | 0.21 (± 0.18; 0.00-0.69) |
| **COMT genotype** | | |
| Val/Val | 7 | 7 |
| Val/Met | 8 | 9 |
| Met/Met | 6 | 5 |

Mean (± standard deviation, range), GSI: Global severity index, SCL-90-R: Symptom-Checklist-90-Revised

* According to the German academic grading system (15-point scale)

** Edinburgh Inventory[1]
